# Origins and Implications of Intron Retention Quantitative Trait Loci in Human Tissues

**DOI:** 10.1101/2024.12.13.628411

**Authors:** Eddie Park, Yi Xing

## Abstract

Intron retention is a type of alternative splicing in which introns remain unspliced in mature RNA transcripts. In order to explore the landscape and consequences of genetically regulated intron retention, we perform an intron retention quantitative trait loci (irQTL) analysis in 49 human tissues across 838 individuals. We identify 8,624 unique intron retention events associated with genetic polymorphisms. 1,369 irQTLs (16%) are also associated with genome-wide association study (GWAS) traits, suggesting that genetically regulated intron retention may play a causal role in the regulation of these GWAS traits. 1,999 irQTLs (23%) colocalize with eQTLs to their respective gene, suggesting that intron retention and steady-state gene expression levels are regulated by shared sets of genetic variants. Through a colocalization analysis of irQTL:eQTL events, we demonstrate that irQTLs are sufficient to generate eQTLs when one of the alternatively spliced transcripts is preferentially targeted by the nonsense mediated decay (NMD) pathway. Surprisingly, for intron retention events whose potential NMD effects can be confidently predicted based on their positions within known gene annotations, we find that 58.8% (923/1570) of the colocalized irQTL and eQTL pairs show effect size directions that are discordant with the NMD model. Moreover, we find that irQTLs are significantly more likely to occur in the same gene with the same effect size direction as compared to exon skipping QTLs. Through mathematical modeling and analysis of experimental perturbation data, we provide evidence that eQTLs are able to generate irQTLs by altering the steady state ratios of spliced and unspliced transcripts, and we postulate that this mechanism may partially underlie the widespread intron retention observed previously in various biological conditions. Taken together, these results show that intron retention and steady state gene expression levels are closely intertwined to regulate phenotypic traits.

## Introduction

Intron retention is a specific class of alternative splicing in which introns are retained in mature RNA transcripts^1,2^. Some early studies have suggested that intronic reads may simply be experimental or transcriptional noise^3^. However, intron retention has been shown to be involved in a wide range of biological processes such as development^4,5^, disease^6,7^, and cancer^8,9^.

Furthermore, intron retention has been suggested to be a source of neoantigens for immunotherapy^10,11^. Intron retention levels (π: PI-Percent Intron inclusion) are measured by the fraction of transcripts containing the intron to the total number of transcripts. Using short read RNA-seq, the levels of spliced and unspliced transcripts are measured by the levels of reads supporting the different isoforms followed by length normalization to account for the unspliced transcript having a greater potential of mapped reads^12,13^.

One of the main consequences of intron retention is to alter transcript stability through nonsense mediated decay (NMD). NMD is a quality control pathway in which mRNAs with premature termination codons (PTC) are degraded^14^. Introns in pre-mRNA are spliced out and exon junction complexes (EJC) remain attached at the spliced sites of processed mRNA. As the mRNA becomes used for translation, ribosomes displace the EJC and prevent NMD. However, if a stop codon prematurely ends translation and prevents the ribosomes from displacing all the EJCs from the RNA, NMD would be triggered, and the transcript would be degraded. The “50- 55 nt rule” for NMD states that NMD would be triggered if the stop codon is located more than 50-55 nt upstream of the last exon-exon^15^. There are many factors that play a role in the efficiency of NMD and the extent of NMD is often difficult to predict^16,17^. If an intron in the coding sequence (CDS) is retained, it may trigger NMD by introducing a PTC within the retained intron or as a consequence of a frameshift to downstream exons. Conversely, if an intron in the 3’UTR is spliced out, this also has the potential to trigger NMD by forming an EJC that is not displaced by translating ribosomes. Alternative splicing (particularly intron retention) coupled to NMD has been described to be an important regulator of gene expression^18^.

In addition to intron retention affecting gene expression, several studies have also demonstrated that changes to gene expression can also have a direct impact on intron retention^19–21^. Using mathematical models and experimental validation, they were able to verify this non-intuitive phenomenon. They were able to show that changing the steady state gene expression level by modulating the transcription rates or degradation rates of RNA transcripts is sufficient to change the steady state intron retention levels of genes. Specifically, in RNA velocity analysis of single-cell RNA-sequencing (scRNA-seq) data, La Manno et al showed that the ratio of unspliced reads to spliced reads can be used to determine whether a gene is being turned on or turned off^19^.

When a gene is turned on, the levels of unspliced reads increase faster than the level of spliced reads. Thus, the measured intron retention levels have a momentary increase in response to increasing transcription. Alternatively, when a gene is turned off, the levels of unspliced reads decrease faster than the level of spliced reads. Thus, the measured intron retention levels have a momentary decrease in response to decreased transcription. This was demonstrated with a simple mathematical model backed by experimental data. However, due to the simplicity of the model, there are several limitations of this model. This model assumes that gene regulation is only controlled by altering the transcription rate of a gene. i.e., the speed at which mRNA transcripts are synthesized. However, biologically, the degradation rate of transcripts also plays an important role in controlling the steady state level of mRNA transcripts^22^. Another limitation of this model assumes a limitless splicing potential in the biological system, which results in the steady state intron retention levels being unaffected by the steady state gene expression levels. It has been documented that, in eukaryotes, the splicing machinery is biologically limited as a mechanism to control the levels of posttranscriptional mRNA processing^23^. Given these limitations of the RNA velocity model, this analysis has been primarily applied under conditions in which the biological system has not reached steady state. By implementing a more sophisticated model, Alkallas et al^20^ and Gaidatzis et al^21^ show that changes to the transcription rate and/or degradation rate of genes have predictable changes to the steady state levels of intronic reads and exonic reads.

Quantitative trait loci (QTL) mapping of molecular traits can be used to identify important regulatory variants for a wide array of molecular phenotypes^24^. For example, expression QTL (eQTL) analysis can be used to identify important cis-regulatory elements that are required for regulating the expression of a particular gene^25^. Splicing QTL (sQTL) analysis has been used to identify important splicing factor binding sites for the efficient splicing of a particular exon^26^.

RNA editing QTL (edQTL) analysis was used to demonstrate that RNA secondary structure plays an important role in the regulation of RNA editing rates^27^.

In this study, we analyzed cis-regulated intron retention events using an intron retention QTL (irQTL) analysis across 49 tissues and 838 individuals from the GTEx project^28^. We find evidence to suggest that eQTLs can be generated from irQTLs by means of altered transcript stability primarily through NMD. Conversely, we demonstrate that irQTLs can also be generated from eQTLs by altering the steady state ratios of spliced and unspliced transcripts.

## Results

### irQTL analysis identifies cis-regulated intron retention events across human tissues

We used an irQTL approach to identify cis-regulated intron retention events and found 8,624 unique introns that are associated with at least one irQTL SNP across 49 tissues (Table S1). Across all tissues, we found introns to be strongly associated with genetic variants (Fig 1a). The number of irQTL events per tissue ranged from 179 in the amygdala to 1,494 in the thyroid (Fig 1b). We observed a correlation (r=0.62) between the number of irQTL events detected and the number of genotyped individuals per tissue. For any given irQTL event, the effect size direction and magnitude were generally consistent across tissues (Fig S1). An example of an intron that is strongly associated with a genetic polymorphism is seen in intron 8 (chr19:762329-7623730) of XAB2 (Fig 1c). Retention of this intron is predicted to trigger NMD by introducing a premature stop codon. XAB2 is described in the literature to have roles in splicing, transcription, and DNA repair^29,30^. Here, a lower intron retention level is associated with the T-allele of rs2335521 while a higher intron retention level is associated with the C-allele (Fig 1d). This is further supported by the T-allele having a stronger MaxEnt splice site score (6.75) while the A-allele had a weaker MaxEnt splice site score (2.99). rs2335521 is within 3 bases of the 5’ splice site of the intron.

**Figure 1.**
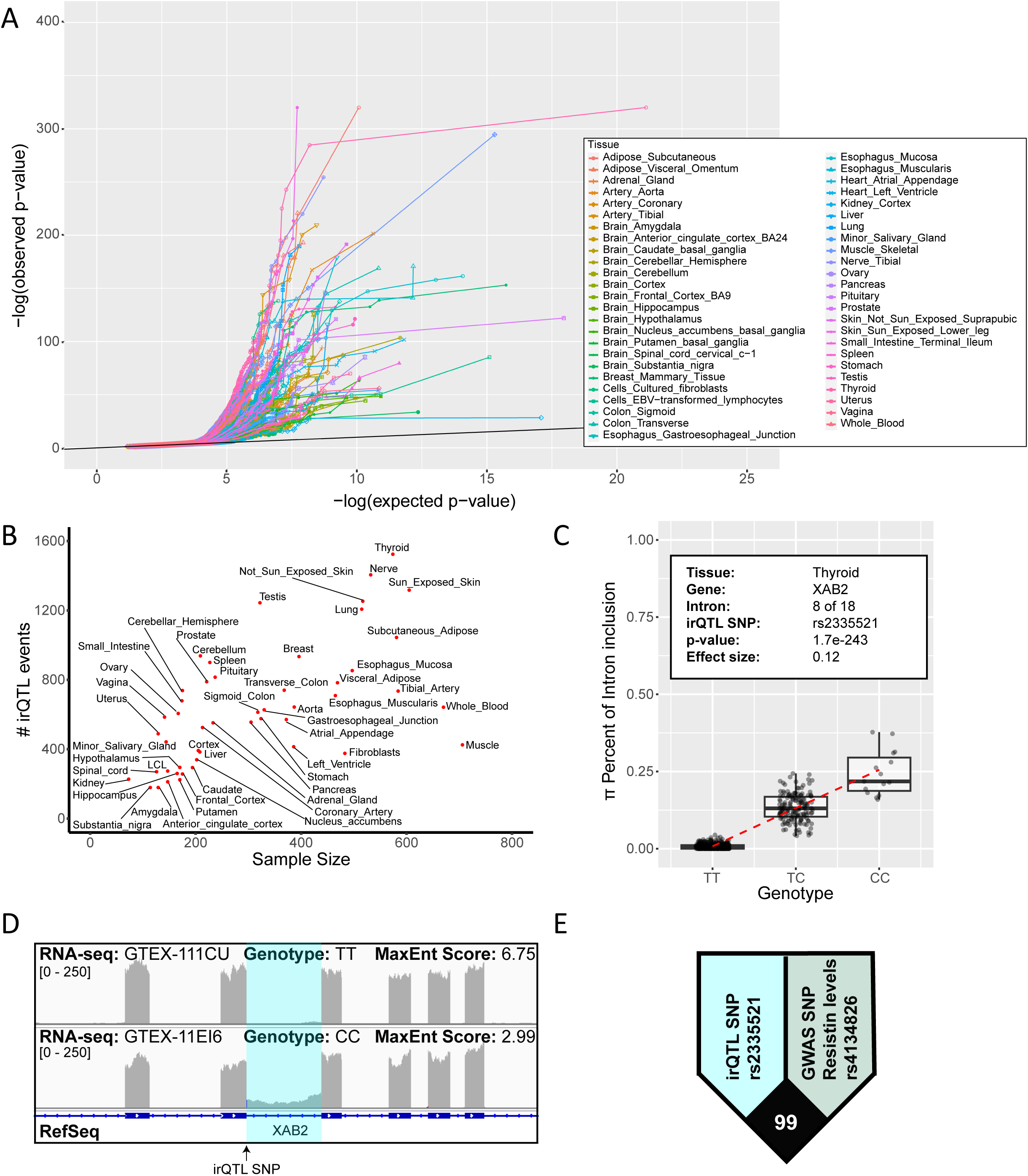
Cis variation of intron retention identified by irQTL analysis. A. Quantile-quantile plot (qq-plot) testing association of intron retention levels with cis genetic polymorphisms across 49 tissues. Black line indicates values for which the observed p value is equal to the expected p-value. B. Scatter plot of the number of irQTL events vs sample size across 49 tissues. C. Example of an irQTL event in the XAB2 gene. Box plots show the significant association of rs2335521 with the intron retention level (π) of intron 8 within the thyroid. Each dot represents data from a particular individual. The dashed red line represents a linear fit of the data. D. An example of two RNA-seq alignments from individuals with homozygous TT and CC genotypes. The irQTL intron is highlighted in light blue. Genome coordinates are flipped to show gene transcription going from left to right. E. LD plot shows GWAS SNP (Resistin levels) linked with irQTL SNP in XAB2.

The irQTL SNP is in high linkage disequilibrium (LD) with a GWAS signal for serum resistin levels (Fig 1e), which suggests that intron retention of intron 8 in XAB2 may result in altered circulating resistin levels in blood. We investigated the extent to which irQTL might be associated with GWAS traits and identified 1,229 unique GWAS traits associated with 1,369 irQTL events (high LD; R2 > 0.8) (Table S2). Taken together, these results suggest that genetic variants have the potential to alter intron retention levels and subsequently influence GWAS traits/diseases.

### Nonsense mediated decay mediates eQTL generation from irQTLs

We hypothesized that one of the main functional consequences of irQTLs would be eQTL generation through NMD of one of the alternatively spliced transcripts. If the irQTL intron is in the CDS, NMD can be triggered from the unspliced transcript if a PTC is introduced as a result of intron retention (Fig 2a). An irQTL coupled with differential transcript stability of one of the transcripts would be sufficient to generate an eQTL for the gene. To identify these types of events, we used colocalization analysis to test whether the irQTL and eQTL are driven by a shared set of genetic variants. Colocalization analysis is a commonly used statistical approach to test whether two QTL signals have shared genetic associations^31^. 1,999 irQTLs colocalized with the eQTL of their respective gene in at least one tissue with a posterior probability of at least 0.75 (Table S3).

**Figure 2.**
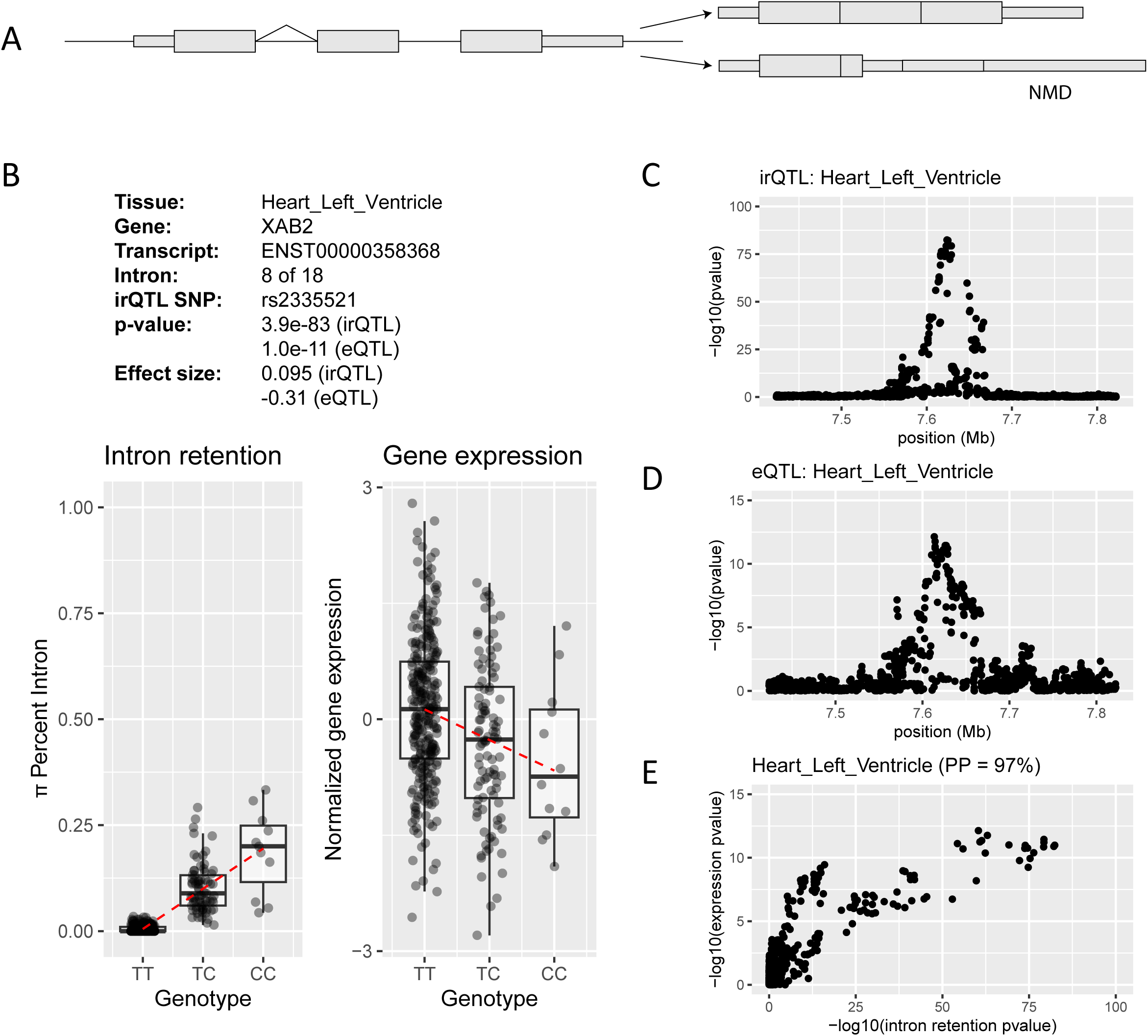
eQTLs generated from irQTLs through NMD. A. Diagram showing how intron retention in the coding region can result in NMD. B. Box plots show the significant association of rs2335521 with the intron retention level (π) at intron 8 and the gene expression level of the XAB2 gene within the left ventricle of the heart. Each dot represents data from a particular individual. Dashed red line represents a linear fit of the data. C. Manhattan plot showing the −log10(p value) for intron retention irQTL in a 400-kb window centered at the intron. D. Manhattan plot showing the −log10(p value) for gene expression eQTL in a 400-kb window centered at the intron. E. Scatter plot of −log10(p value) from irQTL and eQTL signals suggests colocalization of intron retention variation and gene expression variation. Colocalization posterior probability is shown in parentheses.

One example of an eQTL that is likely driven by an NMD-mediated CDS irQTL is in the XAB2 gene (Fig 1cd). The intron retention and gene expression levels are significantly associated with rs2335521 and the irQTL effect size direction is the opposite as the eQTL effect size direction (Fig 2b), consistent with more intron retention leading to more transcript degradation. The Manhattan plots for the irQTL (Fig 2c) and eQTL (Fig 2d) share a similar shape and there is a high colocalization posterior probability (97%) for these two molecular phenotypes (Fig 2e).

Alternatively, when an irQTL intron is in the 3’UTR and sufficiently downstream of the stop codon, the spliced transcript would be subject to NMD and preferentially degraded faster than the unspliced transcript (Figure S2a). One such example of an eQTL that is likely driven by a 3’UTR irQTL is in the FYCO1 gene. FYCO1 is a Rab7 adapter protein and is involved in microtubule transport of autophagosomes^32^. An alternatively spliced intron in the 3’UTR is associated with rs61519998. The T-allele is associated with greater intron retention while the A-allele is associated with greater splicing of the intron (Figure S2b). The intron retention and gene expression levels are significantly associated with rs61519998 and the irQTL effect size direction is the same as the eQTL effect size direction (Figure S2c), consistent with less intron retention leading to more transcript degradation. The Manhattan plots for the irQTL (Figure S2d) and eQTL (Figure S2e) share a similar shape and there is a high colocalization posterior probability (100%) for these two molecular phenotypes (Figure S2f). Taken together, these results suggest that it is possible for eQTLs to be generated from irQTLs if one of the transcripts are targeted by NMD.

### Colocalized irQTL and eQTL pairs are frequently discordant with the NMD model

Surprisingly, within the set of irQTLs that colocalize with eQTLs, there were a few unexpected observations that were unexplainable by the NMD-mediation model described above. When the irQTL intron is in the CDS and introduces a PTC, NMD rules would predict that the direction of the irQTL effect size would be opposite of the eQTL effect size because the unspliced transcript would be predicted to trigger NMD. Conversely, when the irQTL intron is in the 3’UTR, the spliced transcript would be the NMD target. Therefore, the genotype that is associated with lower intron retention should also be the genotype with lower gene expression. Thus, the direction of the irQTL effect size should be the same as the direction of the eQTL effect size.

However, this is often not what we observed in the data. When comparing the direction of irQTL and eQTL effect sizes for introns that are strictly within the CDS in all annotated transcripts of a gene, we observed comparable numbers of irQTL/eQTL pairs with the same effect size directions (875, inconsistent with the NMD model) and opposite effect size directions (637, consistent with the NMD model) (Fig 3a). A similar observation was observed for introns that are strictly in the 3’UTR in all annotated transcripts of a gene (10 same direction, consistent with the NMD model; 48 opposite direction, inconsistent with the NMD model) (Fig 3b).

**Figure 3.**
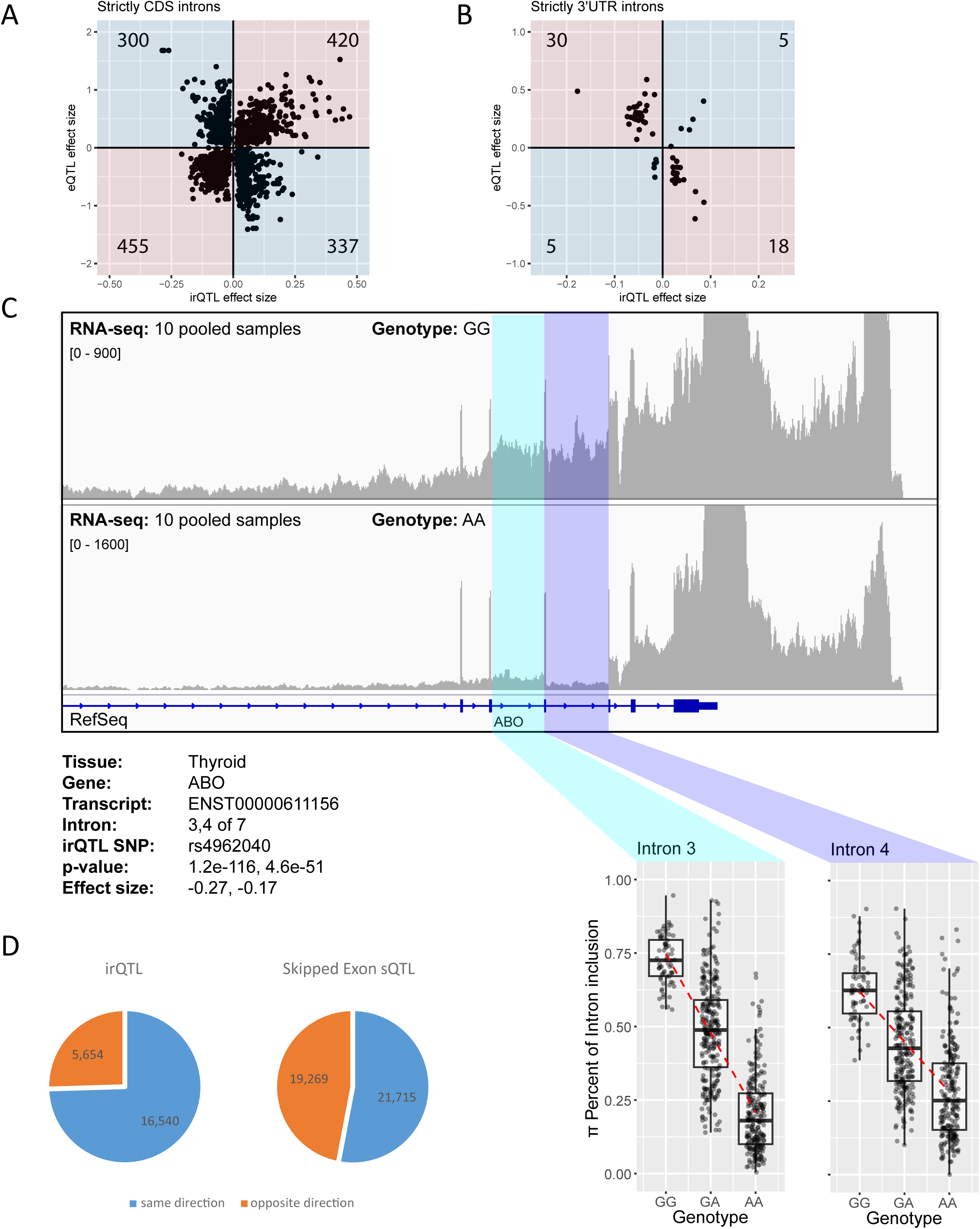
Unexpected observations in irQTL/eQTL analysis. A. Scatter plot of effect sizes in irQTL and eQTL for introns that are strictly in the CDS for all annotated transcripts of a gene. Quadrants concordant with the NMD model are shaded in blue and quadrants discordant with the NMD model are shaded in red. B. Scatter plot of effect sizes in irQTL and eQTL for introns that are strictly in the 3’UTR for all annotated transcripts of a gene. Quadrants concordant with the NMD model are shaded in blue and quadrants discordant with the NMD model are shaded in red. C. Neighboring irQTLs the ABO gene share the same irQTL SNP and have the same effect size direction. Box plots show the significant association of rs4962040 with the intron retention level (π) at intron 3/4 and the gene expression level of the ABO gene within the thyroid. Each dot represents data from a particular individual. Dashed red line represents a linear fit of the data. D. irQTLs in the same gene tend to have the same effect size direction compared to control (skipped exon sQTLs). Pie charts of irQTLs and sQTLs showing whether QTL pairs within the same gene have effect sizes in the same direction or opposite direction.

Another unexpected observation was that many irQTLs had neighboring irQTLs within the same gene. One such example is in the ABO gene. Intron 3 and 4 are both significantly associated with the same set of genetic variants (Fig 3c). Similarly, in this example, other introns within the ABO gene also share the same association. We sought to investigate if this is a widespread phenomenon. We compared the frequency of pairs of irQTLs within the same gene/tissue and compared this to skipped exon sQTL data generated from the same data source as a control. We required the skipped exons to be disjoint to avoid incorrectly double counting the same event.

We found that irQTL pairs within the same genes tend to have effect size directions in the same direction (75%) compared to skipped exon sQTL pairs as control (53%) (Fig 3d) (p-value < 2.2e- 16). This suggests that there may be a common factor that is causing introns within the same gene to have irQTLs at the same time. Furthermore, there may be another mechanism that is contributing to the colocalization between irQTLs and eQTLs. We hypothesized that the gene expression change from an eQTL may be driving multiple irQTLs within the gene.

### Mathematical modeling unifies eQTL and irQTL effects

To explain these observations, we developed a mathematical model to simulate the splicing life cycle of RNA (Fig 4a). Our model is an expanded version of the RNA velocity model^19^. In our model, unspliced RNA (u) is synthesized at a transcription rate (α). We limited the splicing machinery by having a finite amount of spliceosome (E). The spliceosome associates with an unspliced RNA at an association rate (β1) and this unspliced RNA + spliceosome complex (uE) generates a spliced RNA (s) at a splicing reaction rate (β2). Lastly, the unspliced and spliced RNA are degraded at degradation rates (γ1) and (γ2), respectively. An explanation of how the system of differential equations is formulated is provided in the Methods and Supplemental Methods Section.

**Figure 4.**
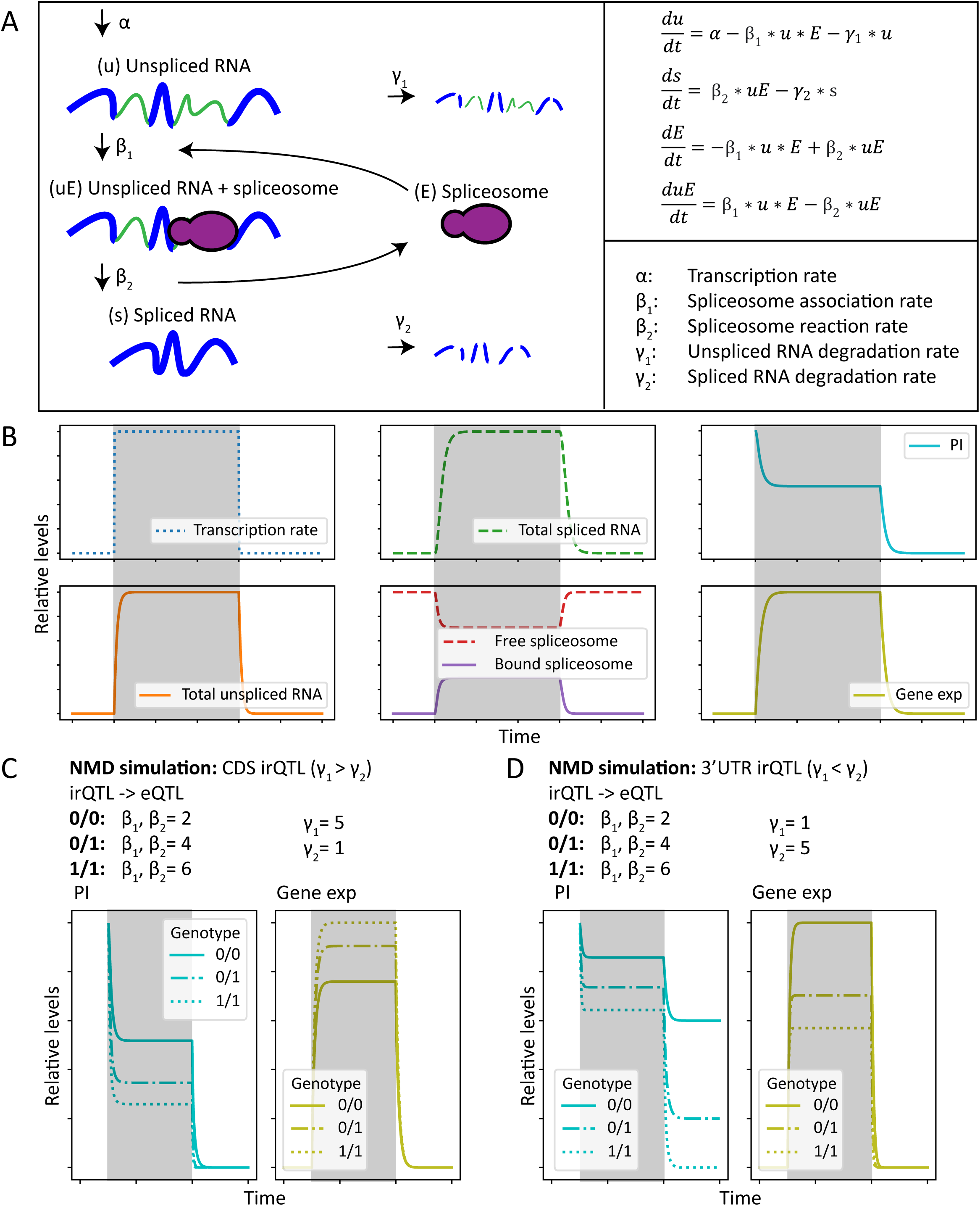
Mathematical model. A. Mathematical model of RNA dynamics. Unspliced RNA (u) is transcribed at transcription rate (α). Unspliced RNA associates with spliceosome (E) at spliceosome association rate (β1) to form an unspliced RNA + spliceosome complex (uE) which is able to generate a spliced RNA (s) at spliceosome reaction rate (β2). Unspliced and spliced RNA are degraded at degradation rates (γ1) and (γ2), respectively. B. Solution of model in (A) in response to step change in transcription (α). Gray boxes represent time intervals in which transcription is turned on. The system is allowed to reach a steady state and then the transcription rate is turned off. C. NMD simulation when the irQTL intron is in the CDS. The degradation rate of the spliced transcript is set to be less than the unspliced transcript. An irQTL is simulated by modulating the spliceosome association rate and the spliceosome reaction rate (β1, β2). D. NMD simulation when the irQTL intron is in the 3’UTR. The degradation rate of the spliced transcript is set to be greater than the unspliced transcript. An irQTL is simulated by modulating the spliceosome association rate and the spliceosome reaction rate (β1, β2).

Our model is equivalent to the RNA velocity model with the following two additions: a limited splicing machinery and allowing the unspliced RNA to degrade without being spliced. Empirical evidence for a limited splicing machinery has been described in literature^23^ and has been used in prior work to model splicing kinetics^20^. The limited splicing machinery contributes to differential intron retention rates in response to differential gene expression at steady state. Allowing the unspliced RNA to degrade was necessary to model the effects of NMD because some forms of NMD are driven by unspliced RNA being degraded without being spliced.

Similar to the analysis used in the RNA velocity study, the solution of our expanded model is solved by setting the transcription rate to be a step function (Fig 4b). Initially, the transcription rate is off. The transcription rate is turned on and the system is allowed to reach a steady state (gray box). Then the transcription rate is turned off. When transcription is on, the total amount of unspliced and spliced RNA reaches steady state levels and subsequently the intron retention (PI) and gene expression levels also reach steady state. Similarly, the spliceosome also reaches a steady state ratio of free and bound levels.

We used this model to simulate the XAB2 example (Fig 1cd, Fig 2) in which eQTLs can be generated from irQTLs when the irQTL intron is in the CDS and the unspliced isoform is degraded at a faster rate by NMD (Fig 4c). In this example, we set the degradation rate of the unspliced isoform to be five times faster than the degradation rate of the spliced isoform and simulated an irQTL by modulating the splicing rate (β1, β2). We simulated three genotypes that had increasing splicing rates (2, 4, and 6). Under these conditions, we observed that the intron retention levels decrease and the gene expression levels increase with increasing splicing rates. Thus, the irQTL and eQTL would have effect size directions in the opposite direction, which is consistent with our empirical observations.

Conversely, we also simulated the opposite case in which the irQTL intron in the 3’UTR triggers NMD so the spliced isoform is degraded faster than the unspliced isoform (Fig 4d). This was the case for FYCO1 (Fig S2). In this situation, through computational simulation, we find the irQTL and eQTL effect sizes are in the same direction, consistent with observations made by RNA-seq.

### eQTLs can generate irQTLs

With this mathematical model, we revisited our earlier hypothesis of eQTLs generating irQTLs. We simulated eQTLs two different ways and investigated the effects on intron retention. Steady state gene expression levels are determined by the transcription rate as well as the degradation rate (Fig 5a). Thus, eQTLs can be generated by genetic variants that alter the transcription rate of a gene or by genetic variants that alter the degradation rate of a gene.

**Figure 5.**
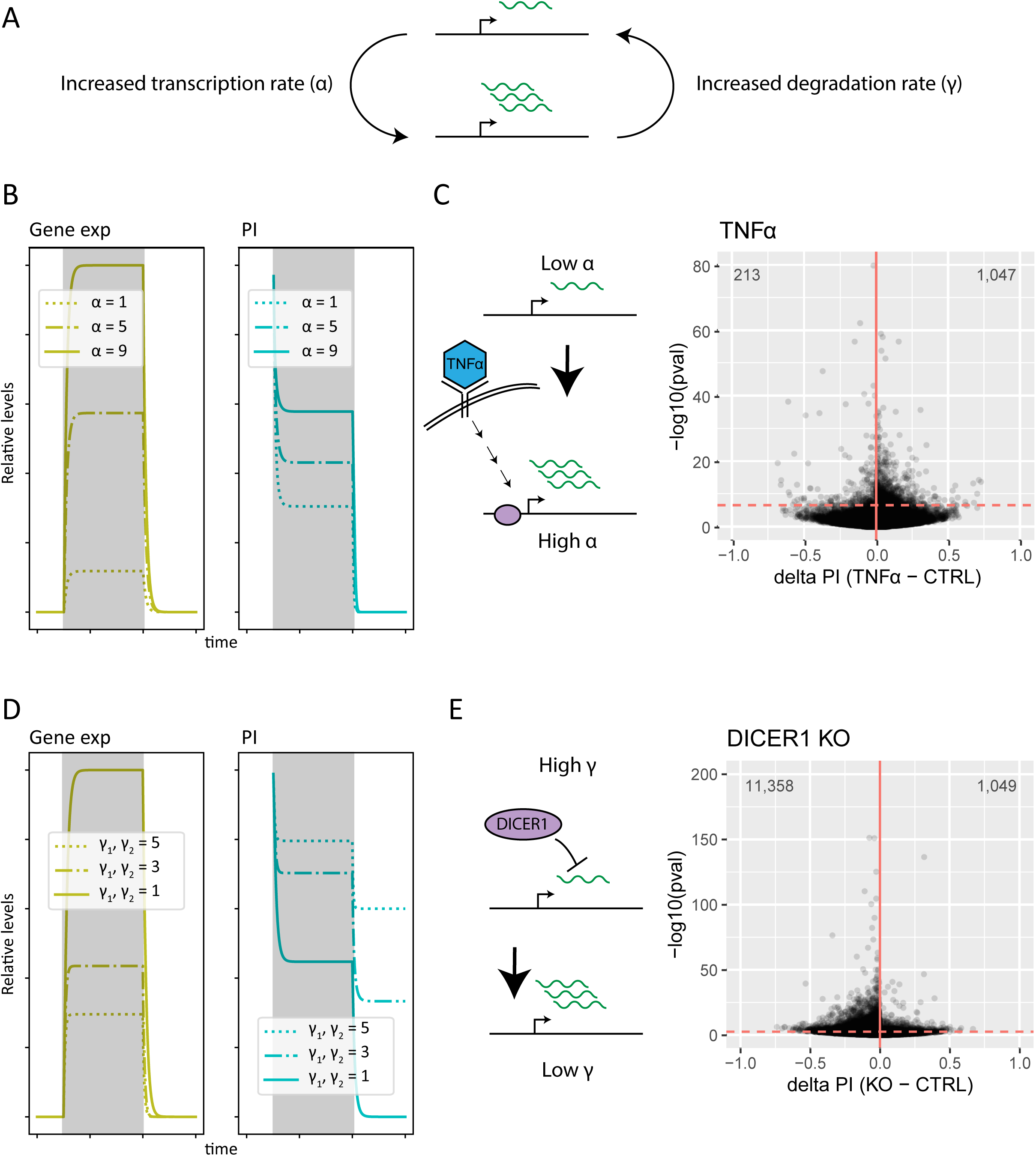
eQTLs can generate irQTLs. A. Steady state transcript levels of a gene are regulated by the transcription rate and the degradation rate. B. Simulation of eQTL generated by changing transcription rate. Left plot shows simulated eQTL. Right plot shows the resulting irQTL. The following parameters were used: β1 = 5, β2 = 5, and E(0) = 1. The other parameters are identical to those described in the Methods. C. Perturbation experiment demonstrating increased intron retention accompanying an increase in the transcription rate. Granulocytic cells are treated with TNF-α. To activate the transcription of TNF-α-induced genes. Dashed horizontal line represents 5% FDR. Numbers in the upper corners represent the number of data points in the upper quadrants of the plot. D. Simulation of eQTL generated by changing degradation rate. Left plot shows simulated eQTL. Right plot shows the resulting irQTL. The following parameters were used: β1 = 2 and β2 = 2. The other parameters are identical to those described in the Methods. E. Perturbation experiment demonstrating decreased intron retention accompanying a decrease in the degradation rate. miRNA processor DICER1 is knocked out in K562 cells. Dashed horizontal line represents 5% FDR. Numbers in the upper corners represent the number of data points in the upper quadrants of the plot.

First, we investigated the effect of altering the transcription rate by increasing the transcription rate (Fig 5b). Using our model, we stimulated an eQTL effect by steadily increasing the transcription rate (1, 5, and 9) of the system while keeping other parameters constant. As expected, in the mathematical simulation, the steady state gene expression level increased with increasing transcription rate. Interestingly, the intron retention level at steady state also increased with increasing transcription rate. We validated this mathematical prediction with a perturbation RNA-seq dataset in which a well-studied signaling pathway is stimulated to activate a set of target genes (Fig 5c). Tumor necrosis factor alpha (TNF-α) is a proinflammatory cytokine that activates the transcription of a large array of genes^33,34^. We investigated the effect that TNF-α- induced gene activation had on intron retention levels by using a dataset in which granulocytic cells were treated with TNF-α^35^. We find that increasing the transcription rate by TNF-α stimulation results in a widespread increase of intron retention levels, which agrees with the results from the *in silico* perturbation of the mathematical model.

Next, we simulated an eQTL caused by a change in transcript stability by steadily decreasing the degradation rate (5, 3, 1) (Fig 5d). Predictably, the model shows that the steady state gene expression levels increase with decreasing degradation rate when other parameters of the model are held constant. Remarkably, the model also predicts that the intron retention levels decrease with decreasing degradation rate. To test this prediction, we used a DICER1 KO dataset^33^ to investigate the effect of decreasing the degradation rate (Fig 5e). DICER1 plays a critical role in miRNA maturation^36^. Thus, knocking out DICER1 would decrease the degradation of a large number of miRNA targets. In this dataset, DICER1 was deleted with CRISPR in K562 cells. We find that reducing RNA degradation rates by reducing DICER1 results in a widespread decrease of intron retention levels, consistent with the predictions of the model.

Taken together, through mathematical modeling and analysis of experimental perturbation data, we find evidence that genetic variants that alter steady state gene expression levels are also able to alter steady state intron retention levels.

## Discussion

In this study, in order to explore the mechanisms that govern intron retention regulation and the functional consequences of these events, we performed an unbiased survey of intron retention events that are genetically regulated in human populations across 49 tissues. Of the 8,624 unique irQTL events, 16% were associated with a GWAS trait and 23% colocalized with their corresponding eQTL.

We show that NMD plays a major role in linking eQTL with irQTL (Fig 6a). When the intron is in the CDS and triggers NMD, the unspliced isoform is degraded faster than the spliced isoform which causes the resulting eQTL to have the opposite effect size direction as the irQTL.

**Figure 6.**
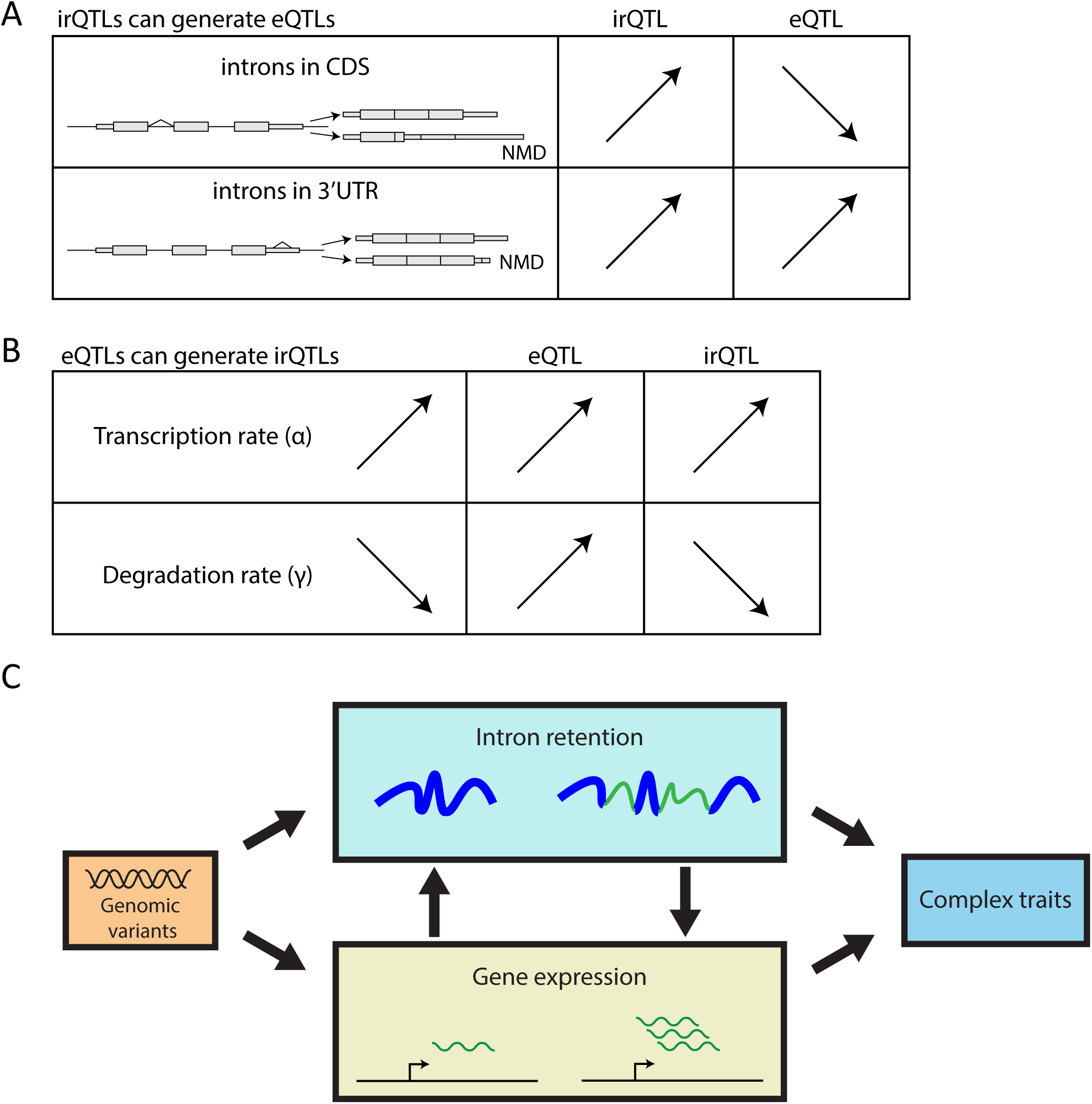
Mutual causation between intron retention and gene expression. A. irQTLs can generate eQTLs through NMD. Arrows indicate effect size direction. B. eQTLs can generate irQTLs through changes in transcription and degradation. Arrows indicate effect size direction. C. Schematic model linking irQTLs, eQTLs and complex traits.

When the intron is in the 3’UTR, the spliced isoform is degraded at a faster rate than the unspliced isoform. Thus, the eQTL that is generated from the irQTL has the same effect size direction as the irQTL.

We had a few unexpected observations that led us to suspect that there may be another mechanism that was playing a role in linking eQTLs with irQTLs beyond NMD. NMD would result in predictable effect size directions for the irQTL and eQTL, based on the location of the intron (Fig 6a). However, we observed effect size directions that did not match expectations (Fig 3ab). Furthermore, we observed that a higher-than-expected number of irQTLs in the same gene have the same effect size direction (Fig 3cd). These observations led us to hypothesize that eQTLs for a gene may be driving multiple irQTLs within the gene, i.e. genetically regulated changes to steady-state gene expression levels could result in genetically regulated intron retention levels. This hypothesis led us to the RNA velocity model^19^, which uses unspliced and spliced reads to predict how RNA profiles will change in a future state. We expanded this model to account for gene expression changes that are driven by RNA degradation as well as RNA synthesis. Furthermore, our expanded model accounts for NMD to show that irQTLs could generate eQTLs if there is uneven degradation of transcript isoforms. The NMD-mediated eQTL generation from irQTL is analogous to results from our earlier work in which eQTL can be generated from RNA editing QTL (edQTL) when one of the transcripts is targeted by miRNA-mediated transcript degradation^37^. Additionally, through the use of mathematical modeling and perturbation experiments, we were able to show that it is possible for irQTLs to arise from eQTLs (Fig 5, Fig 6b). Under conditions in which an eQTL is generated by altering the transcription rate, the irQTL effect size is predicted to have the same effect size direction as the eQTL. Conversely, when the eQTL is generated by altering the degradation rate, the irQTL effect size is predicted to have the opposite effect size direction as the eQTL.

Mathematical modeling provides an important in silico tool to test biological hypotheses to better understand complex biological systems^38,39^. It is a well-established approach to quantitatively explain complex biological phenomena. We were originally led to pursue mathematical modeling to make sense of our data inspired by the RNA velocity model^19^. However, we found that their original model was not appropriate for our analysis (Methods S1), so we made two extensions to the model. The first extension was to limit the enzymatic capabilities of the splicing machinery by modeling a finite amount of the spliceosome. We found that the main effect of this extension is that the steady state levels of intron retention changes with steady state levels of gene expression. Enzymatic limitations to the splicing machinery have been observed to be an important factor in splicing regulation in biological systems^23^ and have also been previously implemented in mathematical models of splicing regulation^20^. The second extension was to allow for unspliced RNA to be degraded without being spliced. The rationale for this extension was so that differences in transcript isoform stability can be modelled to account for NMD in our system.

An important aspect of this work is that intron retention is a special class of RNA processing in which the intron retention level is determined by the fraction of unspliced precursor transcripts over the total amount of unspliced and spliced transcripts. Unlike other types of alternative splicing, intron retention is unique in that one alternative transcript (unspliced) serves as a direct precursor to the other (spliced). Consequently, we expect that other types of alternative splicing, such as exon skipping, would not be affected by this particular type of gene expression driven genetic mechanism (Fig 3d). However, this mechanism could be applied to other types of RNA processing events where one alternative transcript is a direct precursor to the other, such as RNA editing. Thus, a logical implication of this work is that certain biological conditions that induce global changes in gene expression could also drive widespread alterations in intron retention and other RNA processing events. This mechanism may underlie the widespread intron retention and RNA editing observed in various biological conditions^8,40,41^.

In conclusion, we have shown that genetic variants can have direct effects on intron retention and can cause indirect effects on steady state gene expression through alterations in transcript stability by means of NMD. Conversely, we have also shown that genetic variants can have direct effects on gene expression and indirectly alter steady-state intron retention level by changing the flux of RNA metabolism. Previously, it has been shown that splicing QTL (sQTL) can directly affect GWAS traits without affecting gene expression^42^, or indirectly affect GWAS traits by altering steady-state gene expression^43^. Here, our results show that genetically regulated intron retention is closely connected with steady state gene expression and has the potential to alter phenotypic traits (Fig 6c).

## Methods

### Measuring intron retention levels from GTEx RNA-seq datasets

For GTEx V8 samples, RNA-seq alignments (hg38) and genotype information were obtained from the NHGRI’s new genomic data repository Terra platform (https://app.terra.bio/). Intron retention measurements were computed on the cloud using SIRI^12^. We limited our analysis to the 49 tissues with at least 70 genotyped samples. Ensembl 87 was used as the reference gene annotations.

### Preliminary filters of intron retention events for irQTL analysis

For any given tissue, we required introns to meet the following criteria: a minimum average coverage of at least two reads supported the spliced and unspliced versions, a minimum average total coverage of at least ten reads, and a minimum of 10% difference between the intron retention levels of the 90% quantile and the 10% quantile across all individuals. We also removed introns that overlapped with annotated indels from our analysis. Additionally, in order to limit our analysis to bona fide intron retention events, only analyzed U introns as defined by the package SIRI.

### irQTL analysis

For each intron, we applied a linear model to SNPs within a 400-kb window centered at the intron. We used the lm function within R to regress the intron retention level (π) against the genotype across individuals of a given tissue in order to obtain a p value for each SNP. To ensure accurate RNA-seq estimation of intron retention levels, we required each sample to have a minimum coverage of 20 reads. For each SNP, we required a minor allele frequency of at least 5%. For each intron, the irQTL SNP was defined as the closest SNP with the most significant association. We used a p value of 1e-5 as the cutoff to call irQTL events. We defined the irQTL effect size as the slope determined from the linear model such that the y values are the individual intron retention levels (π) and the x values are the genotypes (0, 1, 2 for Ref:Ref, Ref:Alt, and Alt:Alt, respectively).

### GWAS signals

We obtained GWAS signals from the NHGRI-EBI GWAS Catalog^44^ (version: 2021-03-25). VCFtools^45^ was used to calculate linkage disequilibrium (LD) correlations between irQTL SNPs and GWAS SNPs. We required irQTL SNPs to be in high LD (R2 > 0.8) with GWAS SNPs.

Genotypes from the GTEx project were used in the LD calculation. LD plots were generated with Haploview^46^.

### MaxEnt Scores

Splice site scores were computed using MaxEntScan^47^ using genomic sequences with SNP variants.

### Colocalization analysis

We used coloc^31^ for colocalization analysis. GWAS summary statistics were obtained from the NHGRI-EBI GWAS catalog^44^. irQTL summary statistics were generated with a linear model described above. eQTL summary statistics were obtained from the GTEx Portal^28^.

### Mathematical modeling

The transcription rate (α) is modelled as a set function going from 0, 1, and back to 0, unless otherwise specified. The unspliced RNA can associated with the spliceosome at a rate of β1*u*E to form a complex with the spliceosome (uE). The unspliced RNA can also be directly degraded at a rate of γ1*u. (uE) is created from (s) and (E) at a rate of β1*u*E and the splicing reaction reform (E) and creates a new spliced RNA (s) at rate β2*uE. Lastly, the spliced RNA (s) is degraded at rate γ2*s. The mathematical models were solved numerically using the scipy.integrate subpackage^48^. The parameters we used were α = 1, β_1_ = 1, β_2_ = 1, E(0) = 2, γ_1_ = 1, γ_2_ = 1, unless otherwise specified. Here, we have explored the parameter space thoroughly and find that the qualitative trends are preserved for a wide range of parameters. But we have selected a set of parameters that best show the desired outcomes.

### Perturbation experiments

RNA-seq data were obtained from TNF-α^35^, DICER KO^33^, and Actinomycin D^49^ datasets were obtained from publicly available sources. Reads were aligned to hg38 using STAR version 2.7.0d^50^. Replicates were merged and SIRI was used to measure intron retention levels. Ensembl 87 was used as the reference gene annotations. p-values were calculated using a Fisher’s exact test.

## Acknowledgements

The Genotype-Tissue Expression (GTEx) Project was supported by the Common Fund of the Office of the Director of the National Institutes of Health, and by the NCI, NHGRI, NHLBI, NIDA, NIMH, and NINDS.

---

**Supplemental Figure 1**

Heatmap of irQTL effect sizes for 138 introns (rows) across 49 tissues (columns). Only introns with sufficient coverage to pass filters and have effect sizes computed across all 49 tissues are displayed in the plot.

**Supplemental Figure 2**

A. Diagram showing how intron retention in the 3’UTR can result in nonsense mediated decay.

B. An example of an intron retention event in the 3’UTR of FYCO1. Two RNA-seq alignments from individuals with homozygous TT and AA genotypes are shown. The irQTL intron is highlighted in light blue. Genome coordinates are flipped to show gene transcription going from left to right.

C. Box plots show the significant association of rs61519998 with the intron retention level (π) at intron 6 and the gene expression level of the FYCO1 gene within the tibial artery. Each dot represents data from a particular individual. Dashed red line represents a linear fit of the data.

D. Manhattan plot showing the −log10(p value) for intron retention irQTL in a 400-kb window centered at the intron.

E. Manhattan plot showing the −log10(p value) for gene expression eQTL in a 400-kb window centered at the intron.

F. Scatter plot of −log10(p value) from irQTL and eQTL signals suggests colocalization of intron retention variation and gene expression variation. Colocalization posterior probability is shown in parentheses.

**Supplemental Table 1**

List of irQTL events across 49 tissues.

**Supplemental Table 2**

List of irQTL events associated with GWAS traits (R2 > 0.8).

**Supplemental Table 3**

List of irQTL events that colocalize with eQTL (PP4 > 0.75).

