## Supplementary material for "Origins and Implications of Intron Retention Quantitative Trait Loci in Human Tissues": MethodsS1.docx

**Methods S1. Expanded description of mathematical modeling**

Eddie Park, Yi Xing

**The need for an expanded RNA velocity model**

Our mathematical modeling analysis was originally inspired by the model from the RNA velocity study^1^ (Fig 1a). We started with this model to help explain some of the non-intuitive observations from our data. However, we came across a few issues that showed that the original RNA velocity model from La Manno et al was inadequate for our analysis. So we made an expanded version for our analysis.

In the original RNA velocity model, a simple mathematical model is established to describe the relation between changing gene expression (controlled at the level of transcription) and changing ratio of unspliced reads to spliced reads. Here, the transcription rate (α) is set as a step function: starting in the off state, then turning on for a period of time before turning off. Unspliced RNA (u) is synthesized and as the unspliced RNA accumulates it is converted to spliced RNA (s) at a splicing rate (β). The spliced RNA is degraded at a degradation rate (γ). We defined the intron retention levels (PI: Percent Intron) to be the ratio of unspliced RNA to the total levels of RNA. We define gene expression levels to be the total levels of RNA in the system (Fig 1b).

Even though the RNA velocity model was initially established for scRNA-seq analysis, we found that it can be readily applied to bulk RNA-seq analysis. We simulated the effect of turning off transcription once the system has reached a steady state. The gene expression levels are expected to decrease once the transcription rate is turned off, but the RNA velocity model also shows that the intron retention levels also decrease (Fig 1c). This is also verified with experimental data in which 293T cells at steady state are treated with actinomycin D (a transcriptional inhibitor) and a global decrease in intron retention levels are observed^2^ (Fig 1d).

Although the RNA velocity model captures dynamic changes to intron retention in response to changes to the transcription rate, the model fails to capture intron retention changes at steady state. We simulated three different transcription rates and observed an expected change to steady state gene expression levels, but the intron retention level remained identical at these different transcription rates (Fig 1e). Furthermore, NMD cannot be simulated from the RNA velocity model because unspliced RNA is not permitted to degrade without being spliced before degradation.

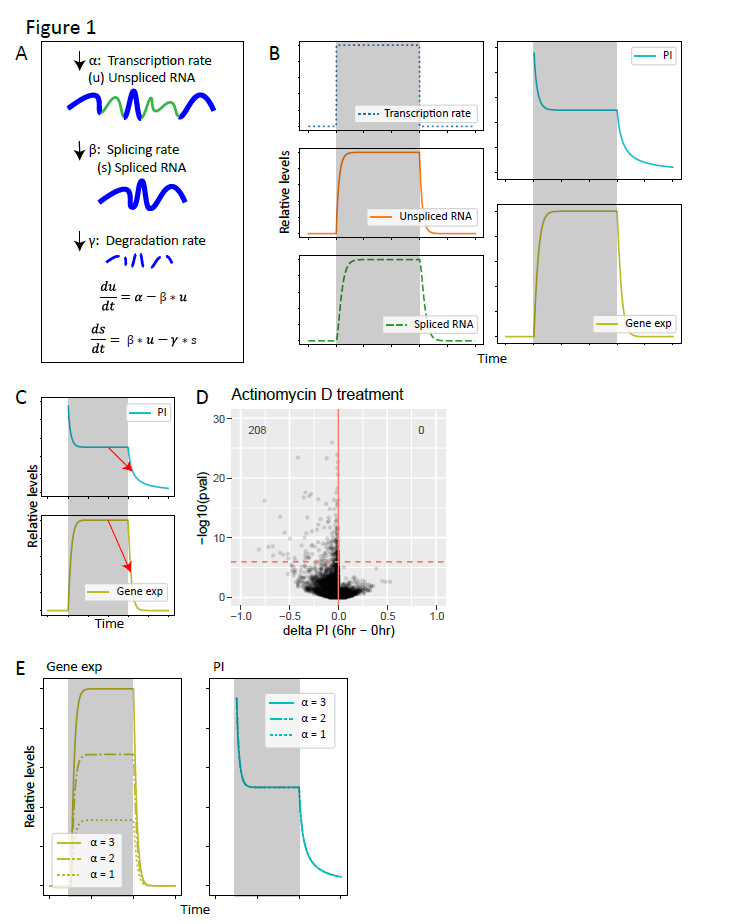

**Figure 1**

A.	Original RNA velocity model. Unspliced RNA (u) is transcribed at transcription rate (α). Unspliced RNA is spliced at rate (β) to form spliced RNA (s). Spliced RNA is degraded at rates (γ).

B.	Solution of model in (A) in response to step change in transcription (α). Gray boxes represent time intervals in which transcription is turned on. The system is allowed to reach a steady state and then the transcription rate is turned off.

C.	Simulating the effect of turning off transcription. Red arrows indicate the expected change in intron retention and gene expression in response to turning off transcription. The system is at a resting steady state and the transcription rate (α) is set to zero.

D.	Experimental data showing the effect of turning off transcription. Actinomycin D is used to turn off transcription in 293T cells. Dashed horizontal line represents 5% FDR. Numbers in the upper corners represent the number of data points in the upper quadrants of the plot.

E.	Simulating increasing a transcription rate. The effects on gene expression and intron retention are shown.
