## Supplementary figures and images for "Origins and Implications of Intron Retention Quantitative Trait Loci in Human Tissues"

### FigureS1.pdf

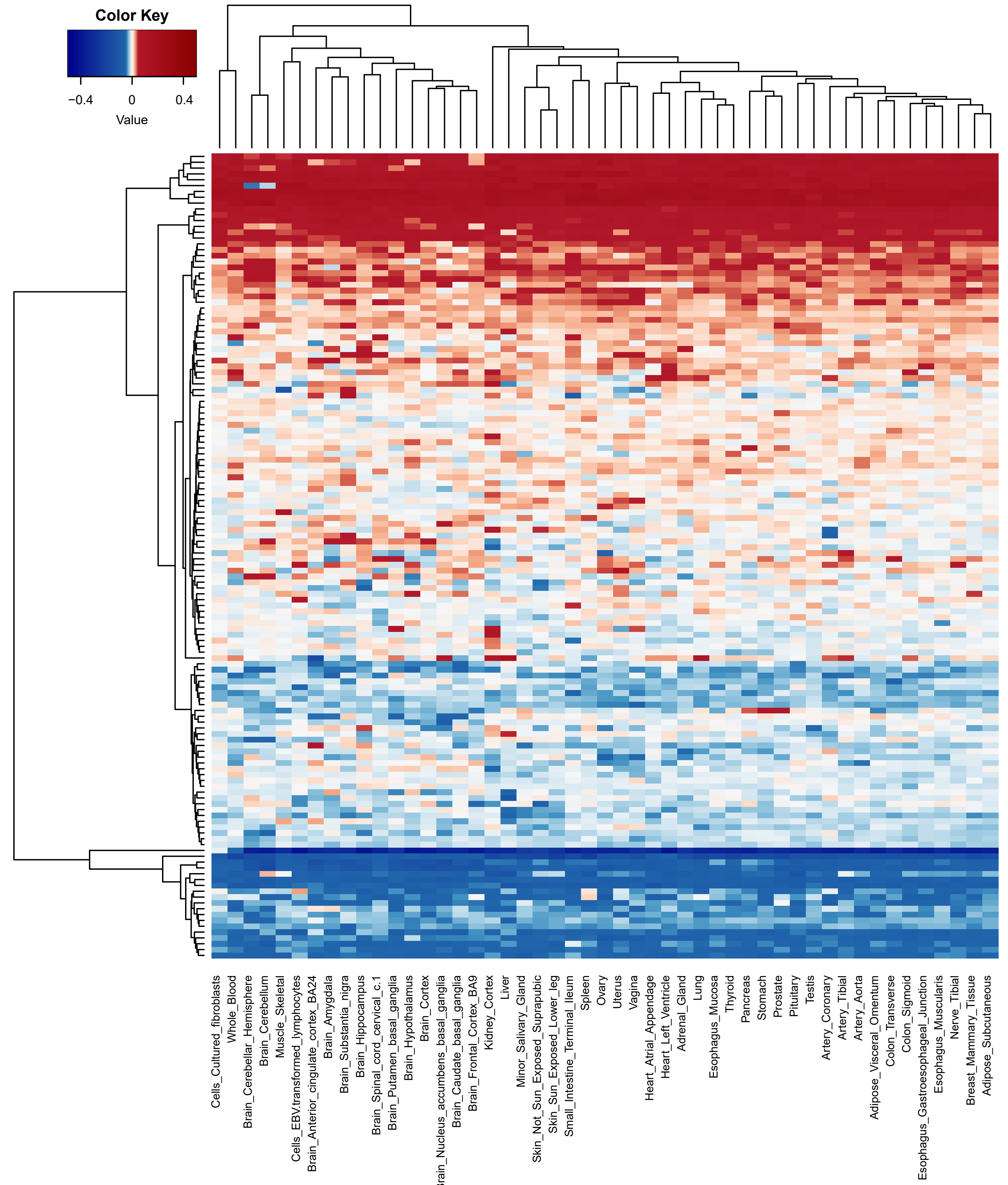

### FigureS2.pdf

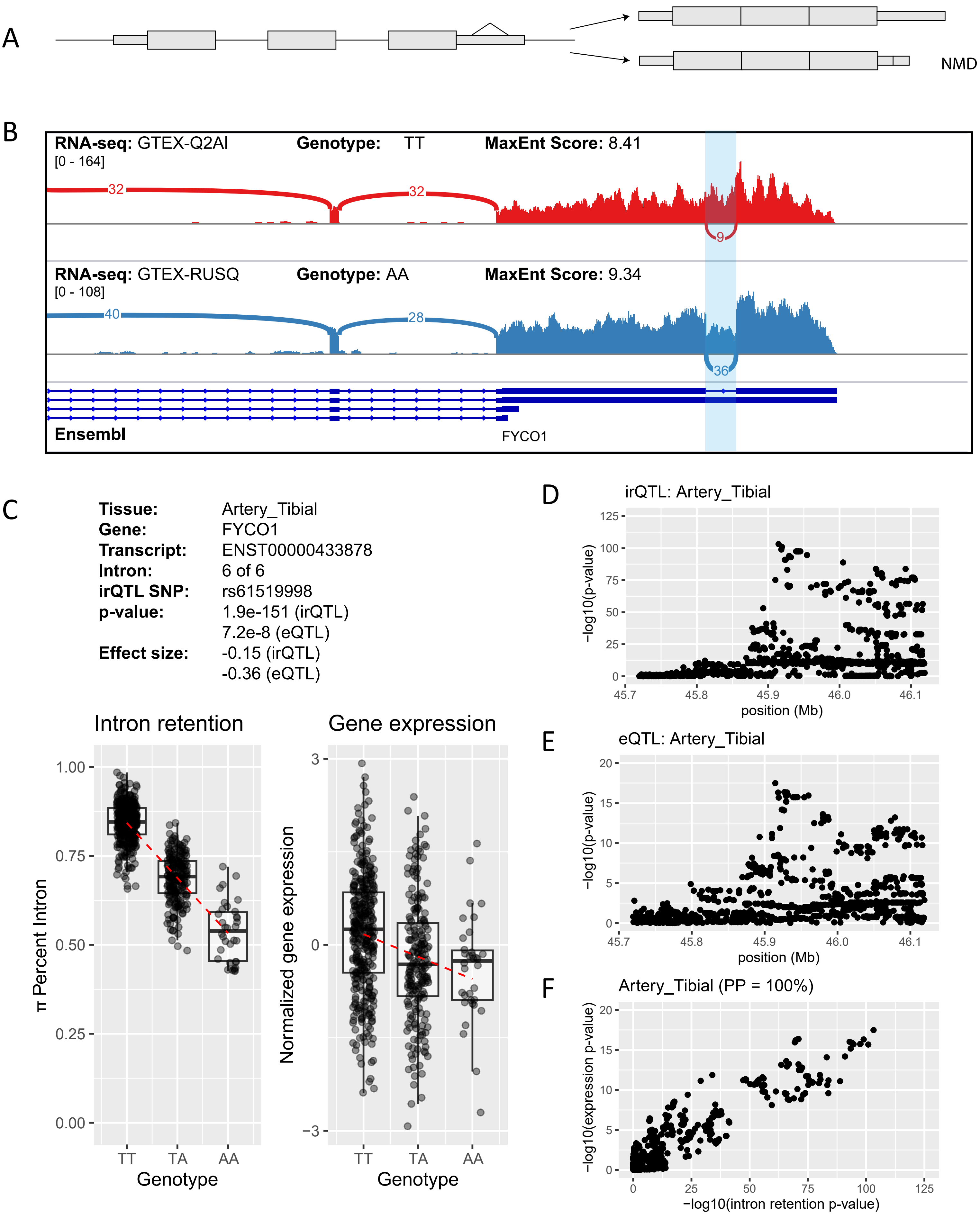
